# NaviCom: A web application to create interactive molecular network portraits using multi-level omics data

**DOI:** 10.1101/089367

**Authors:** Mathurin Dorel, Emmanuel Barillot, Andrei Zinovyev, Inna Kuperstein

**Author notes:** Last co-authors, or (Andrei Zinovyev, Inna Kuperstein).

## Abstract

Human diseases such as cancer are routinely characterized by high-throughput molecular technologies, and multi-level omics data are accumulated in public databases at increasing rate. Retrieval and visualization of these data in the context of molecular network maps can provide insights into the pattern of molecular functions encompassed by an omics profile. In order to make this task easy, we developed NaviCom, a Python package and web platform for visualization of multi-level omics data on top of biological network maps. NaviCom is bridging the gap between cBioPortal, the most used resource of large-scale cancer omics data and NaviCell, a data visualization web service that contains several molecular network map collections. NaviCom proposes several standardized modes of data display on top of molecular network maps, allowing to address specific biological questions. We illustrate how users can easily create interactive network-based cancer molecular portraits via NaviCom web interface using the maps of Atlas of Cancer Signaling Network (ACSN) and other maps. Analysis of these molecular portraits can help in formulating a scientific hypothesis on the molecular mechanisms deregulated in the studied disease.

## Introduction

Today’s biology is largely data-driven, thanks to high-throughput technologies that allow investigating molecular and cellular aspects of life at large scale. These technologies comprise microarray platforms, next-generation sequencers, mass spectrometers, or interaction screens, producing multi-level omics data such as gene and protein expression, mutational profiles, epigenetic landscapes, etc (1, 2). Samples of diseased tissues, especially in cancer, are routinely characterized by these high-throughput techniques, and multi-level omics data are accumulated in public and proprietary databases. More than 20 000 tumor samples have been profiled so far by two main international efforts, The Cancer Genome Atlas (TCGA, <u>www.cancergenome.nih.gov</u>) and the International Cancer Genome Consortium (ICGC, <u>www.icgc.org</u>). Multiple cancer omics datasets are available through the Gene Expression Omnibus (GEO, <u>www.ncbi.nlm.nih.gov/geo</u>) and the cBioPortal web interface (<u>www.cbioportal.org</u>) (3). An integrated analysis of different data types can provide the most comprehensive picture on the status of the studied sample: however this represents a challenge because of the complex cross-correlations in the structure of multi-level omics data (4). Data visualization is a possible solution to obtain an integrated overview of the data and understanding their characteristic patterns. Making biological sense out of the molecular data requires visualizing them in the context of cell signaling processes (5, (6). A lot of information about molecular mechanisms is available in the scientific literature, and is also integrated into signaling pathway databases. Those signaling pathway databases, such as Reactome (7), KEGG PATHWAYS (8), Spike (9), PathwaysCommons (10), ConsensusPathDB (11), SIGNOR (12), Atlas of Cancer Signaling Network (ACSN) (13) etc, vary in the approach to depict molecular interactions and in the level of details of biological processes representation (14). Several data visualization tools address the problem of exploiting omics data in the context of biological pathways and networks. These tools, such as Ipath (15), BioCyc (16), Pathway projector (17), several plugins in Cytoscape (18), 19) or NaviCell (20) can be used together with a number of pathway databases. In addition, some pathways databases contain integrated data visualization tools such as the Reactome Analysis tools or FuncTree for KEGG PATHWAYS (21).

Nevertheless, there remain several poorly addressed problems in the field. First, exploiting network maps and pathway databases for the visualization of multi-level omics data faces the challenge of compatibility between the databases, data types and visualization methods. Users without computational biology background may face difficulties when choosing the most appropriate tool for their purposes. The second bottleneck lies in accessing public omics data resources and retrieving the data, for instance in order to compare the original results obtained by a research group with publicly available molecular profiles. There are currently no tools that support automated import of large datasets from omics data resources and displaying them on top of molecular network maps with optimized visualization settings. The third challenge is that there is no standard graphics representation to visualize multiple omics data at the same time that would provide to the user the best combination of data visualization channels.

To fill these gaps, we developed NaviCom, a tool for automatically fetching and displaying several omics data types on top of network maps, using optimized pre-defined data visualization modes. NaviCom connects cBioPortal to NaviCell and allows visualizing various high-throughput data types simultaneously on a network map. To highlight the possibilities provided by this tool, we demonstrate how multi-level cancer omics data from cBioPortal (3) are automatically visualized on the molecular network maps available in ACSN (13) and NaviCell (22) collections.

Integrating together different type of data allows to create the comprehensive molecular portraits of cancer, identifying specific patterns in the data that may lead to better disease subtypes classification (23, 24). However, the ‘classical’ definition of molecular portraits does not consider signalling networks. NaviCom allows the user to create complex interactive molecular portraits of cancers based on simultaneous integration and analysis of multi-level omics data in the context of signalling networks. This broaden the possibilities of data interpretation, allowing not only to pattern the traits of changes in the data, but also to understand involvement of specific molecular mechanisms associated with the studied disease.

## Materials and Methods

### Data source and type

We use available omics data from cBioPortal (3) (<u>www.cbioportal.org</u>), a web resource for the exploration of cancer genomic datasets from TCGA project and several other projects. The studies on the cBioPortal database contain large-scale cancer data sets including expression data for mRNA, microRNA, proteins; mutation, gene copy number, methylation profiles and beyond. cBioPortal data can be extracted per gene or per patient using the R package cgdsr, an R connection to the Cancer Genomic Data Server API, a REST-based programming interface.

Datasets used for application examples described below are: Breast Invasive Carcinoma (TCGA, Nature 2012), Acute Myeloid Leukemia (AML) (TCGA, NEJM 2013), Adenocortical Carcinoma (TCGA Provisional), Ovarian Serous Cystadenocarcinoma (TCGA, Nature 2011), Glioblastoma (TCGA, Cell 2013), Sarcoma (TCGA, Provisional)

### Signaling network maps

Any map prepared in NaviCell format can be used for visualization of data via NaviCom. To demonstrate this feature, we provide three visualization examples on maps with different characteristics: i). The Atlas of Cancer Signaling Network (ACSN, <u>https://acsn.curie.fr</u>) (13) that contains a comprehensive description of cancer-related mechanisms retrieved from the recent literature, following the hallmarks of cancer (25). Construction and update of Atlas of Cancer of Signaling Networks (ACSN) is done using CellDesigner tool (26), involving manual mining of molecular biology literature. ii). Alzheimer’s map (27) created using Cell Designer tool (26). iii). Ewing’s Sarcoma signaling map (28) created in map drawing tool of Cytoscape. Last two maps are part of the NaviCell collection (<u>https://navicell.curie.fr/pages/maps.html</u>) (22).

### NaviCom software design

NaviCom is a web interface to the python package navicom and the R package cBioFetchR to import, format and display data from cBioPortal on the signalling maps using NaviCell visualization procedures (Figure 1). The interaction with cBioPortal is performed using our R package cBioFetchR, an easy to use wrapper around the R package cgdsr. The interaction with NaviCell is managed by our python module navicom, which defines optimized visualisation modes for various types of high-throughput data. The list of studies available on cBioPortal is generated dynamically when the NaviCom web page is opened, thus presenting up-to-date information to the user. NaviCom also extracts the number of samples, and the types of data available in each dataset. Once the visualisation is requested, NaviCom accesses a cached copy of the dataset, or downloads it directly from cBioPortal with cBioFetcher to generate this local copy. The data are then loaded in NaviCell by the navicom module according to the visualization mode that is requested through NaviCom. NaviCom provides a default visualization setting (detailed below) for simultaneous integration of the data into the big comprehensive maps of molecular interactions. Once chosen, these settings are applied automatically, significantly reducing the time required to perform the visualization comparing to manual mode. It also allows launching the visualization of several datasets on different maps in parallel. In addition, since the whole dataset is already imported to NaviCell in a form of data tables, the user also may apply different types of analyses provided by the NaviCell environment. Finally, the tool can be run using the command line that opens further possibilities for data analysis (Supplementary table 1).

**Figure 1.**
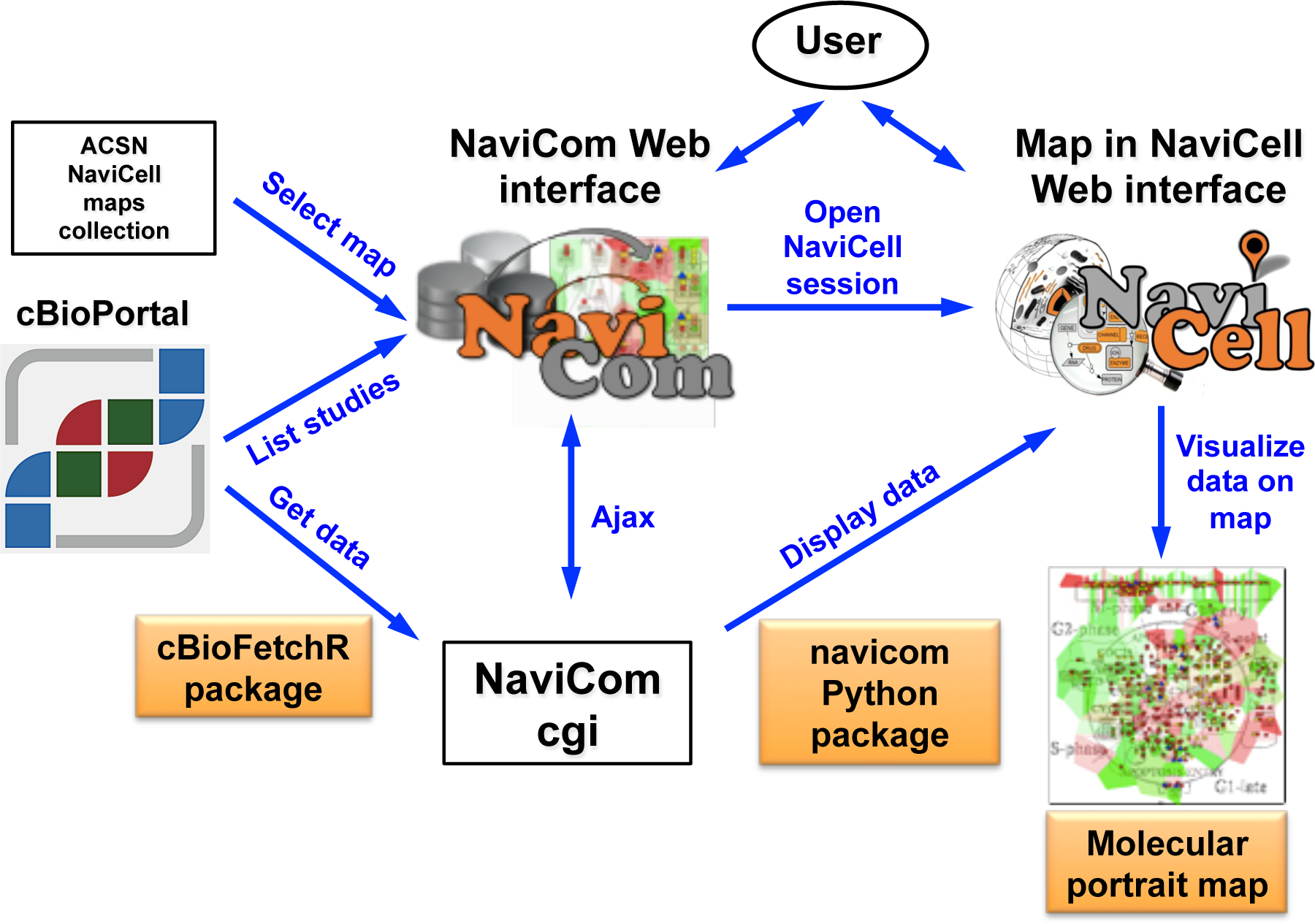
General architecture of NaviCom. The NaviCom interface provides the user with an updated list of studies from cBioPortal and links to ACSN and NaviCell maps collections. When visualization is requested, NaviCom starts a new NaviCell session and calls a cgi on the server. The cgi downloads cBioPortal data to the NaviCell session and displays them to generate the molecular portrait selected by the user.

### NaviCom data visualization

Navicom uses combinations of the visualisations channels available in NaviCell. Those multiple channels offer different ways displaying the data on top of the network map, which allow the user to visualize several layers of information simultaneously on one map in a comprehensive manner.

Data representation through multiple channels correspond to various graphics:

> **Barplots** are histograms located near each entity, with the height and color of the bar depending on the value of the data for the entity.
>
> **Heatmaps** consist in plotting colored squares near each node, with the color depending on the value of the data for the node.
>
> **Glyphs** consist of geometrical figures with variable shape, color and size depending on the values of the data for the entity.
>
> **Map staining** is a unique display mode for coloring background areas around each entity according to the data value associated to this entity.

### Navicom display settings

NaviCom uses combination of the NaviCell graphics described above, to overlay multiple data types. To avoid time-consuming manual manipulation, the visualization settings are standardized as pre-defined data display modes (Table 1). These NaviCom settings are applied automatically, making the visualization process easy for users, also allowing launching visualization of several datasets on different maps in parallel. The visualization is applied to the entire dataset, and the user can then choose subgroups using the interactive interface of NaviCell.

There are four major pre-defined NaviCom display modes:

> **Complete data type display mode** overlays as many information as possible on the map in order to compare the coherence of the various signals, and to spot the most affected areas on the signaling map. This mode is suitable for exploratory analysis and comparison of samples.
>
> **Triple data type display mode (Mutations and genomic data)** shows genomic and transcriptomic data together: mRNA expression as map staining, copy number variation as heat map and the mutations as glyph, together allowing the understanding the alterations of each gene at different regulatory levels.
>
> **Double data type display mode** shows each one of the available data type in the context of mRNA expression as map staining, which allows comparison of the profile in the displayed omics (e.g. methylation) data with the transcriptional status of the gene.
>
> **Single data type display mode** is the simplest and fastest visualization mode allowing user to focus on distribution of one type of data over the map.

In addition, a user can choose to “Export the dataset to NaviCell”, which will grant access to all the datatables of the dataset with a preconfigured display settings on a NaviCell map, to perform different analysis of the data.

**Table 1.**
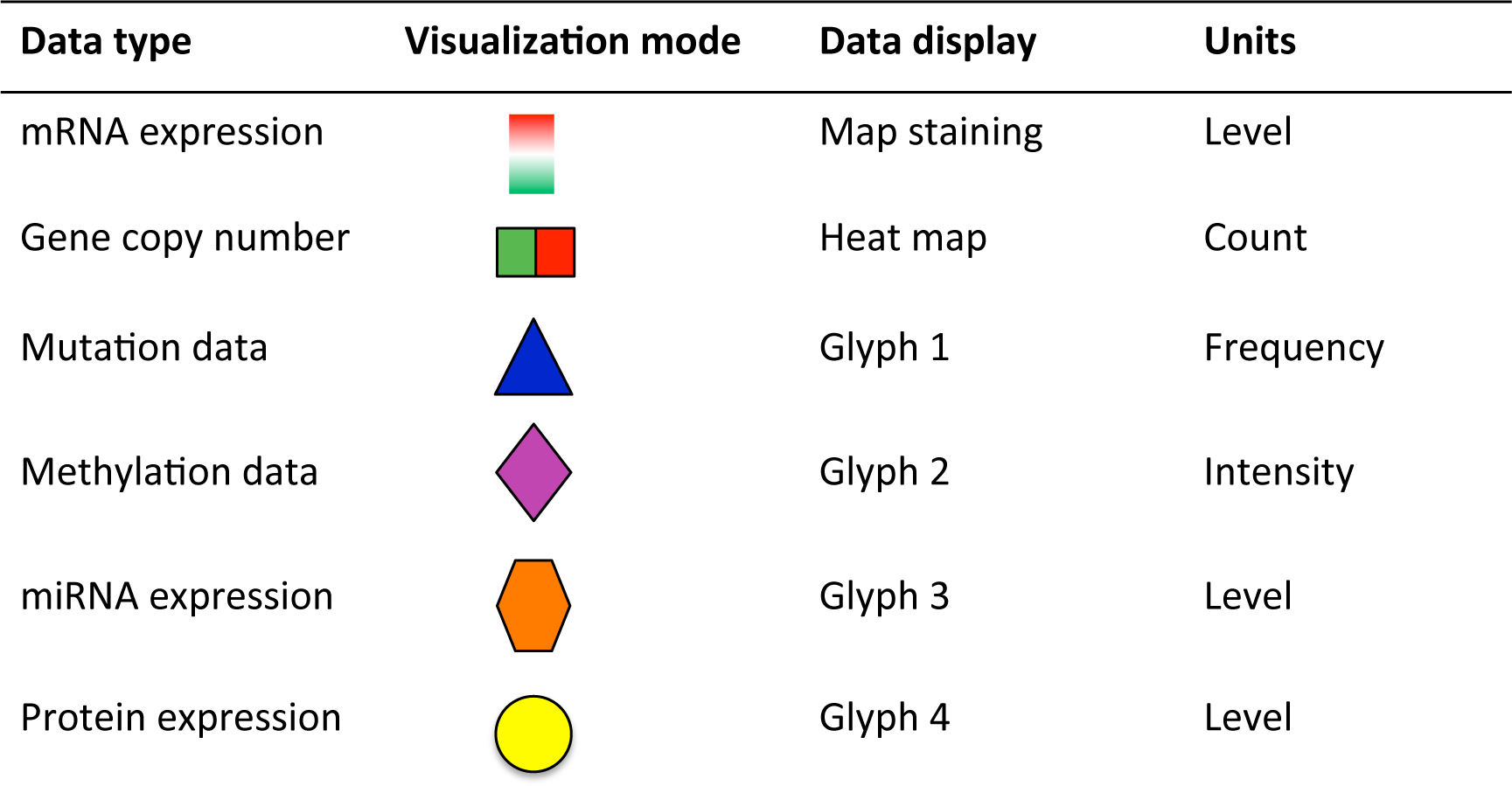
Data display settings in NaviCom

| Data type | Visualization mode | Data display | Units |
| --- | --- | --- | --- |
| mRNA expression |  | Map staining | Level |
| Gene copy number |  | Heat map | Count |
| Mutation data |  | Glyph 1 | Frequency |
| Methylation data |  | Glyph 2 | Intensity |
| miRNA expression |  | Glyph 3 | Level |
| Protein expression |  | Glyph 4 | Level |

### NaviCom access, availability and documentation

The NaviCom web page is freely accessible via <u>http://navicom.curie.fr</u>. cBioFetchR and *navicom* codes are available at <u>https://github.com/sysbio-curie/cBioFetchR</u> and <u>https://github.com/sysbio-curie/navicom</u> respectively. NaviCom, cBioFetchR and *navicom* are distributed under LGPL license. A detailed documentation including introduction into the NaviCom features, code description, manual, tutorial and application examples are available at <u>https://navicom.curie.fr/downloads</u>.

## Results

We demonstrate a new tool NaviCom that allows to optimize usability and increases compatibility between existing resources. We show how NaviCom solves the problem of connecting the omics data resource cBioPortal and the collection of signaling maps in ACSN and NaviCell; and optimizes data visualization.

We first provide an overview of NaviCom workflow, thus describe the features of NaviCom comparing to other efforts in the field and finalize by NaviCom application examples, including molecular portraits of several cancer types.

### NaviCom workflow

The web-based user interface for NaviCom makes it easy to display omics data from cBioPortal on interactive maps provided in NaviCell format (<u>http://navicom.curie.fr</u>). The workflow consists of several simple steps (Figure 2): **i) Data**: selection of the dataset from the list available in cBioPortal. A short summary (types of data and number of samples available) is available for each datasets. **ii) Map**: selection of the molecular network map from ACSN or NaviCell collections to visualize the data on. The user can import and visualize data via NaviCom on any type of signaling networks. **iii) Display modes**: selection of the data visualization procedure may vary depending on the data available in the data set and the scientific question. **iv) (optional). Display configuration**: the data display default settings are pre-defined by NaviCom (Table 1): however users may adjust the color gradient settings. Once chosen, these settings are applied on the NaviCell map, significantly reducing the time required to perform the visualization comparing to manual mode. The map with data displayed according to the chosen configurations will be opened in a separate tab containing a NaviCell session.

**Figure 2.**
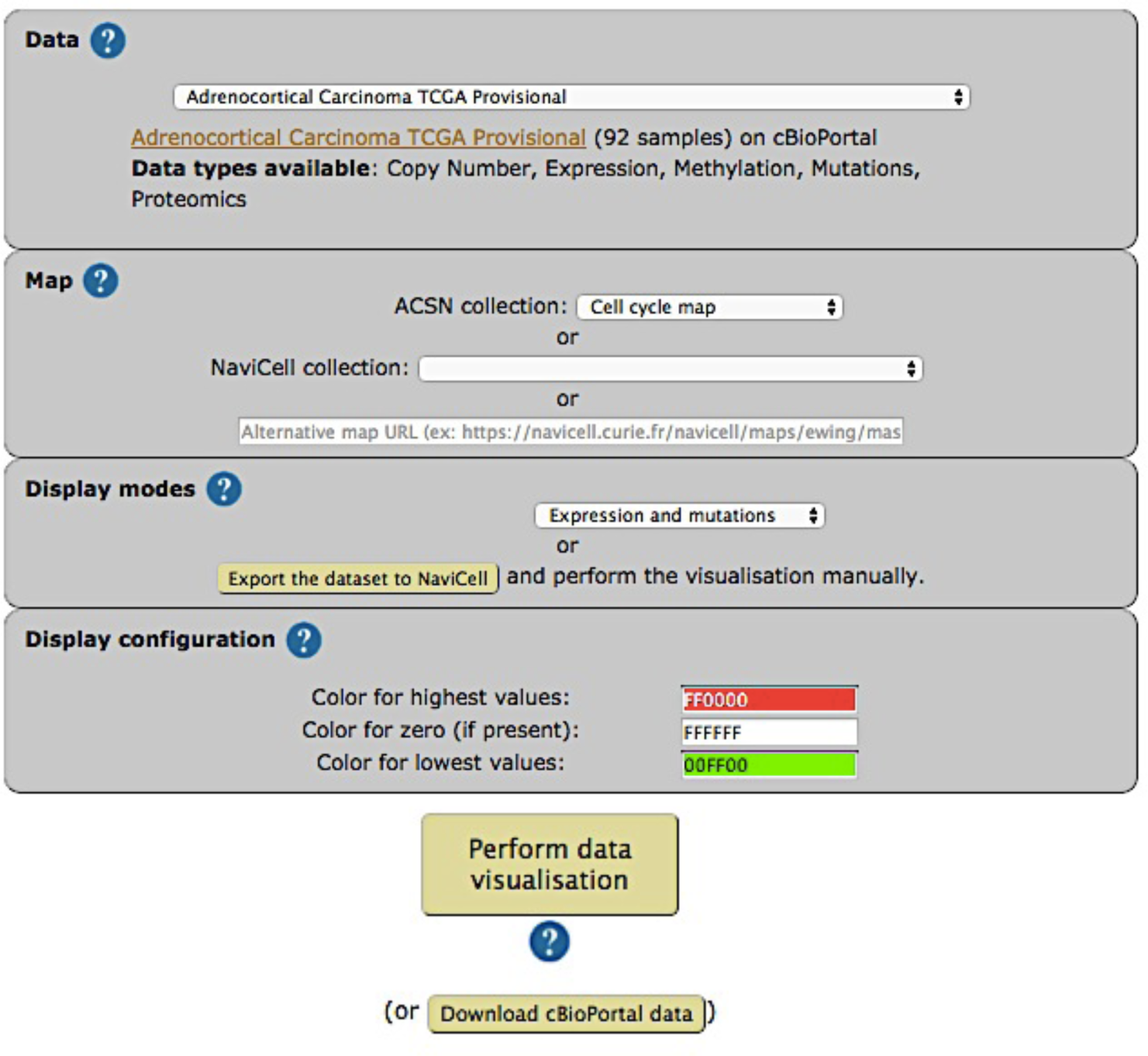
Visualization setting panel of NaviCom.

The resulting maps with visualized data on top of them are interactive and can be browsed using NaviCell Google Maps-based navigation features (Figure 3). For example, semantic zooming allows the user to interact with the maps starting from the top level view (Figure 3A), where patterns of integrated data can be grasped, up to the most detailed view at the level of individual actors (Figure 3B, C).

**Figure 3.**
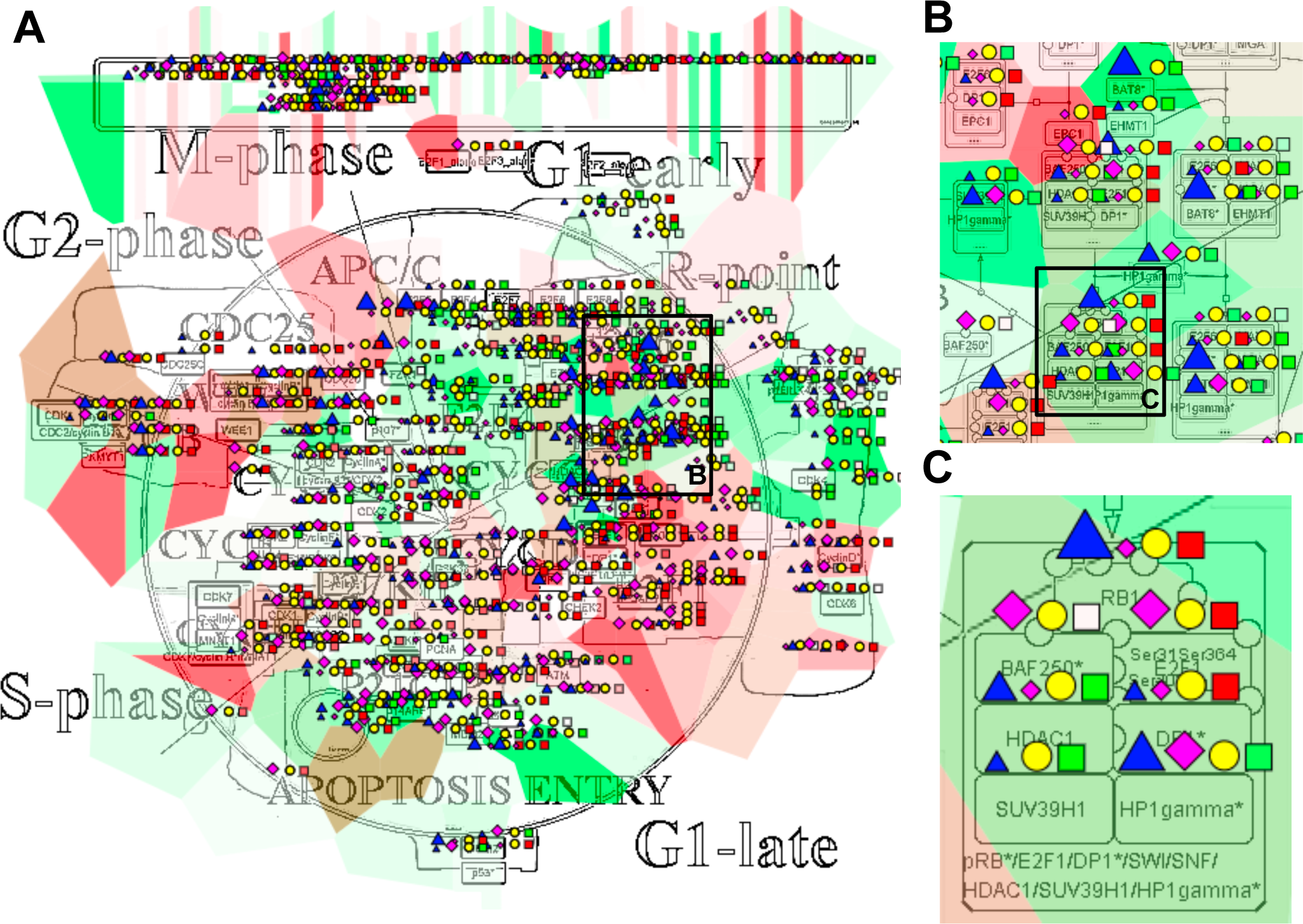
Multi-omics data visualization in Cell Cycle signaling map. Five types of omics data, Copy Number, Expression, Methylation, Mutations, Proteomics, for Breast Invasive Carcinoma dataset from cBioPortal has been displayed on the map using the pre-defined display mode as detailed in Table 1. The values represent average for 825 samples available in the dataset (A) Top level view of data distribution, (B) and (C) Zoom in on individual entities on the map.

### NaviCom features

NaviCom is characterized by the combination of the following essential and unique features: i) Accessibility via a user-friendly web interface; ii) Fully integrated tool allowing automatic fetching and display of various omics data types on molecular network maps; iii) It offers several meaningful scenarios of multi-level omics data visualization. The data pre-defined display modes in NaviCom are optimized for simultaneous visualization of several data types, which makes the visualization process easy for users; iv). The user can import and visualize data via NaviCom on any NaviCell map. NaviCom allows exploiting not only pre-defined but also user-created pathway maps; v) The tool can be run using the command line, allowing the integration into data analysis pipelines with little efforts; vi) The resulting ‘colored’ interactive maps with displayed data are characteristic complex molecular portraits of cancer types.

### Comparison of NaviCom to similar tools

We compared NaviCom functionalities to similar tools available in the field (Table 2). For example, Cytoscape (18) is a widely used network visualization software which provides several plugins to analyse data in a network context, and some to perform simple visualization (CyLineUpi, PINA4MSi, CytoHiC), but does not allow overlay of several type of data. The BiNoM (29) plugin currently provides two types of data display: coloring of individual map nodes and map staining using pre-defined ‘territories’ of functional modules for displaying average values of expression for all genes in each module. However, BiNoM requires external CellDesigner screenshots, and does not provide many visualization modes. FuncTree (30) is a web application for analysis and visualization of large scale genomic data. It is linked to KEGG PATHWAYS (8) and offers many functionalities like normalization or enrichment analysis. It can display data as circles of varying diameter, but does not allow the display of several types of data at the same time and requires a lot of interactions from the user. iPath (31) is a web-based tool for the visualization of data on pathways maps where the size and color of the nodes can be set to match various data, which allows a very clear visualization. However, iPath requires the colors and width to be specified as inputs, and provides no processing functionalities. It also lacks a convenient zooming and provides visualization on metabolic maps only. Pathway Projector (32) is a web-based pathway browser which allows custom data visualization. It can use KEGG PATHWAYS maps and provides many modes of display for the data. However, this power comes at the cost of a very complex interface, with highly descriptive input files, which require heavy preprocessing of the data. Reactome (33) is a pathway database which provide mapping of data on its pathway maps. It is very useful, by its completeness, for enrichment analysis, but does not allow the visualization of the values of the data. BioCyc (16) is a collection of pathway databases similar to Reactome which provides tools for visualizing multi-omics data on pathway map. However this visualization is limited to a large scale view of the pathway, which hinders the ability to interpret such visualization for large scale data, and only provides a complex REST API.

**Table 2.**
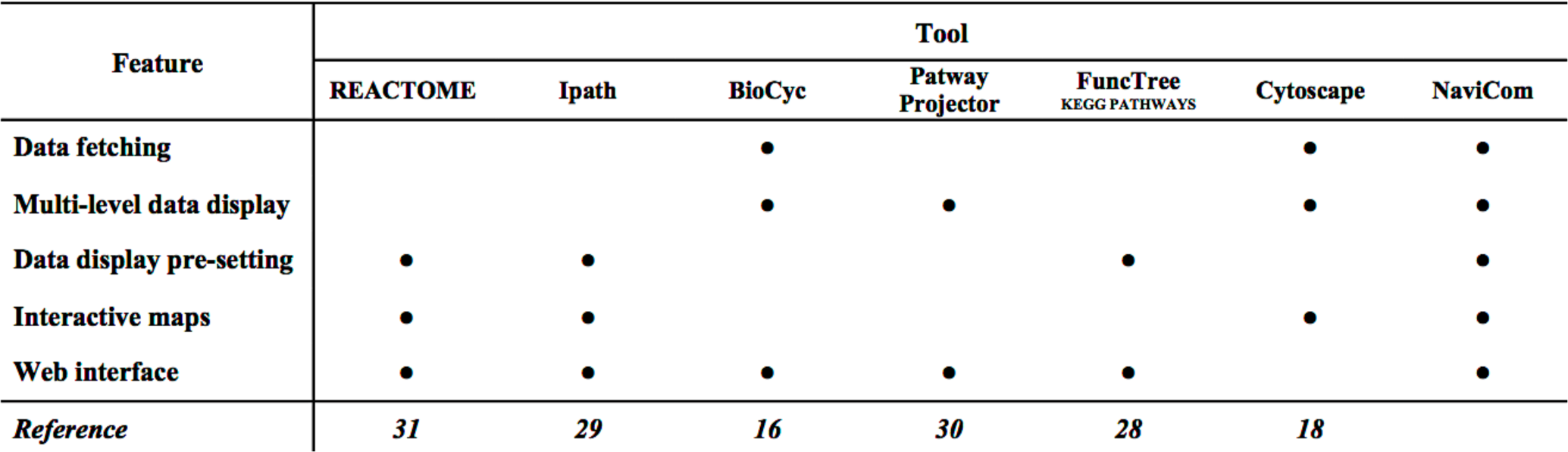
Comparison of NaviCom with similar tools

The most important advantages of NaviCom web application over existing similar tools is that it combines several essential functions together: allows fetching the data and visualizing several large multi-level omics dataset from cBioPortal on top of comprehensive molecular maps even for inexperienced users. The combination of these features is missing in the currently existing visualization tools.

### Molecular portraits of cancer

We define a molecular portrait as a visualization of molecular traits of a biological sample or a group of samples on top of a signaling map. In the first example we use the Cell Cycle map from the ACSN collection, created in CellDesigner and presented in the NaviCell format. Figure 3 shows the map with the complete display of the Breast Invasive Carcinoma dataset, consisting of 5 type of omics data. This type of display provides the most comprehensive picture of data, observed at the top level view (Figure 3A). The map with displayed data is interactive, data distribution on individual entities can be observed while zooming in (Figures 3 B and C). In the second example the Alzheimer’s disease signaling map from NaviCell collection, that is also created in CellDesigner pathway editor, is used to visualize three types of data (expression, copy number and mutations) from the Glioblastoma dataset (Supplementary figure 1). In addition, any biological network map that can be imported into Cytoscape environment can be also used for data display using NaviCom, as it is shown for the Ewing’s Sarcoma signaling map, demonstrated with the complete display of the Sarcoma dataset (Supplementary Figure 1).

Since NaviCom uses aggregated values for all samples in each dataset from cBioPortal, it helps to generate interactive molecular portraits of cancers, facilitating comparison between cancer types. This is demonstrated in Figure 4, where the DNA repair map from ACSN collection is colored by expression and mutation data from four different types of cancers. The supplementary figure 3 shows more extensive visualization of the same data, using the whole ACSN map. Such comparisons allows to grasp the patterns in data across molecular mechanisms in a comprehensive manner, helping to highlight the specific deregulated processes in each type of cancer and highlight and deduce deregulated ‘hot area’ specific to each type of cancer. These signaling network-based molecular portraits and signatures that can be derived from them are of help for patients’ stratification.

**Figure 4.**
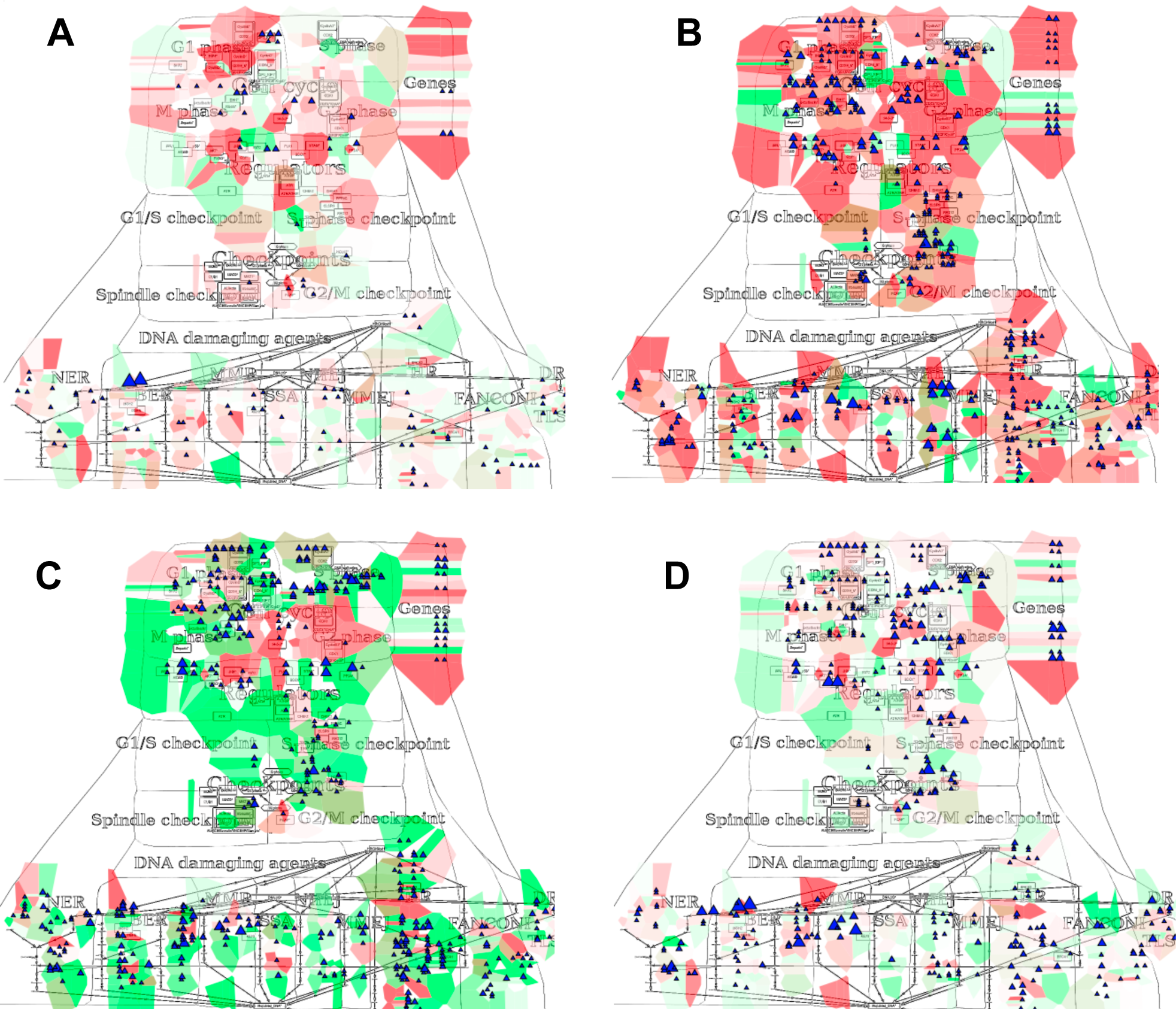
Molecular portraits of cancer types. Expression and mutation data from CbioPortal has been displayed on the DNA repair map using the pre-defined display mode as detailed in Table 1. The values represent average for all samples available in each dataset (A) Acute Myeloid Leukemia, (B) Adenocortical Carcinoma (C) Ovarian Serous Cystadenocarcinoma, (D) Glioblastoma. The molecular portrait demonstrate clear difference between the four types of cancer. Notable that Adenocortical Carcinoma data set is characterized by massive activation of cell cycle modules and high number of mutations, whereas the over three types of cancer are most probably less proliferative, as the cell cycle does not show up and over-activated. Remarkable that Acute Myeloid Leukemia dataset shows the least mutation frequencies.

## Discussion

Combining together different types of high-throughput data provides a more complete description of alterations in a given condition, such as the set of genetic, epigenetic and post-translational alterations in cancer. Data visualization is a powerful method allowing quick grasp of the main points of interest in high-throughput data. These molecular descriptions placed in the context of biological processes depicted in a form of interactive maps, orient the researcher in identifying the deregulated mechanisms in studied samples and diseases. Signaling maps contain information about the connectivity between biological entities, therefore data visualization performed on top of network maps provides the possibility to retrieve entities relationship information to understand changes in molecular processes under different conditions.

Researchers in biology (and cancer biology) are often faced with the need to access public database resources for comparing their results with public molecular profiles (e.g. of tumor) and for analysing molecular observations in the context of signaling network map. Various omics data are available on public and local databases (4). However, there are no tools that support import of big datasets from these databases and displaying them on signaling network maps in efficient way and with optimized visualization settings. To answer to this demand, we developed NaviCom for automatic simultaneous display of multi-level data in the context of signaling network map, provided in a user-friendly manner.

We envisage several directions in further development of NaviCom. We plan to include network-based data processing methods such as geographical smoothing (over the neighbors in the network) to account for the extra information provided by the network. Similarly, NaviCom could provide pre-grouping of the data per various features available in cBioPortal, such as the grade of the disease, the age the patient, the survival status and generate directly a gallery of molecular portraits for various subtypes. So far, NaviCom is optimized for one omics data resource to be linked to one type of signaling maps collection (cBioPortal to ACSN/NaviCell maps collections). However NaviCom design can be used to bridge any type of data and networks resources. In the near future we will extend the NaviCom platform to provide access to a wide range of omics databases (starting with ICGC, HGMB, CCLE etc.). In addition, in order to allow a broader description of the molecular mechanisms implicated in the studied samples, signaling networks available in databases such as KEGG PATHWAYS (34), Reactome (35) SIGNOR (12) and others, will also be integrated and used for multi-level omics data analysis via NaviCom.

The platform will facilitate investigation of various biological conditions, tissue/cell types and diseases. For instance, NaviCom allows fast visualization of patients’ data in the context of maps and comparison with existing portraits from the associated gallery of molecular portraits of cancers different cancer types and sub-types available at the NaviCom website, helping for patient stratification.

## Acknowledgements

We thank Eric Bonnet for technical advices and help in website generation. We thank Maria Kondratova for testing NaviCom. This work has been supported by the Agilent Thought Leader Award #3273; ASSET European Union Framework Program 7 project under grant agreement FP7-HEALTH-2010-259348; grant ‘Projet Incitatif et Collaboratif Computational Systems Biology Approach for Cancer’ from Institut Curie. This work has received support under the program «Investissements d’Avenir» launched by the French Government and implemented by ANR with the references ANR-11-BINF-0001 (Project ABS4NGS) and APLIGOOGLE program provided by CNRS. Funding for open access charge: Institutional budget of institut Curie - INSERM U900 department.

## Supplementary materials

**Supplementary table 1.**
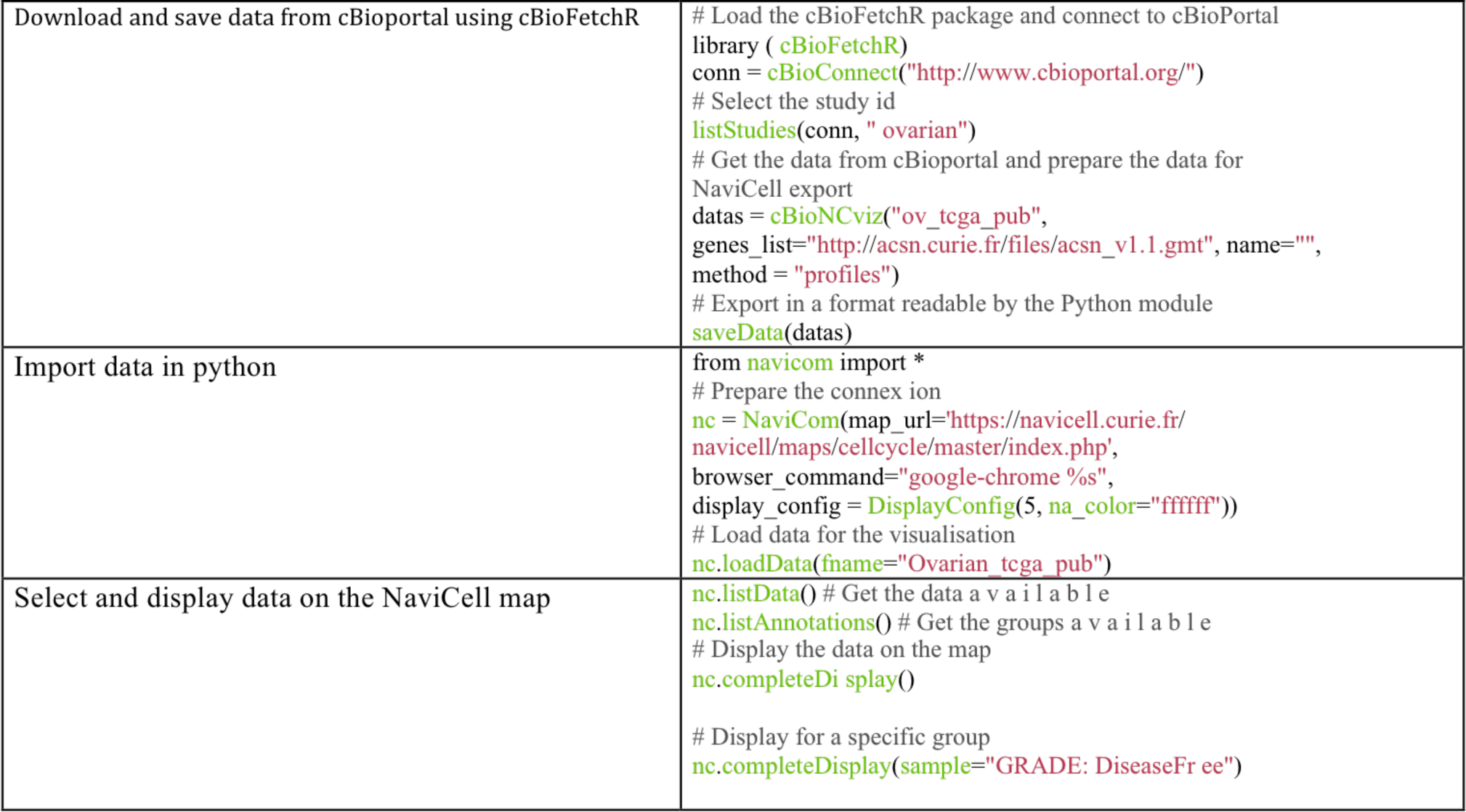
Using NaviCom from the command line. The activity of NaviCom requires the R package cBioFetchR and the Python module navicom. With those packages, it is possible use NaviCom with the command line, granting more flexibility in various ways, such as a more precise configuration of the display in NaviCell or the list of genes downloaded from cBioPortal.

**Supplementary Figure 1.**
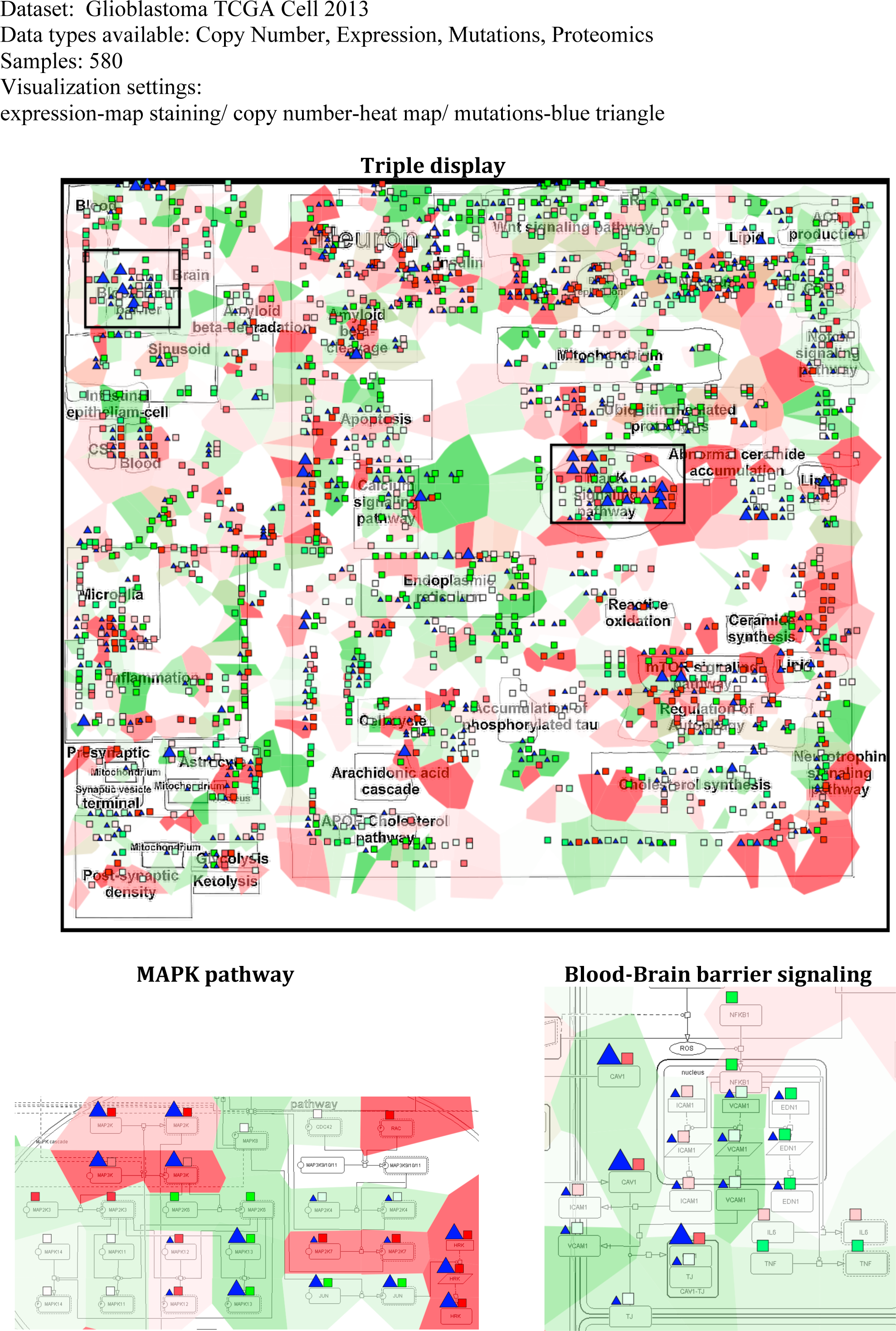
Molecular portrait of Glioblastoma visualized on Alzheimer’s signaling map.

**Supplementary figure 2.**
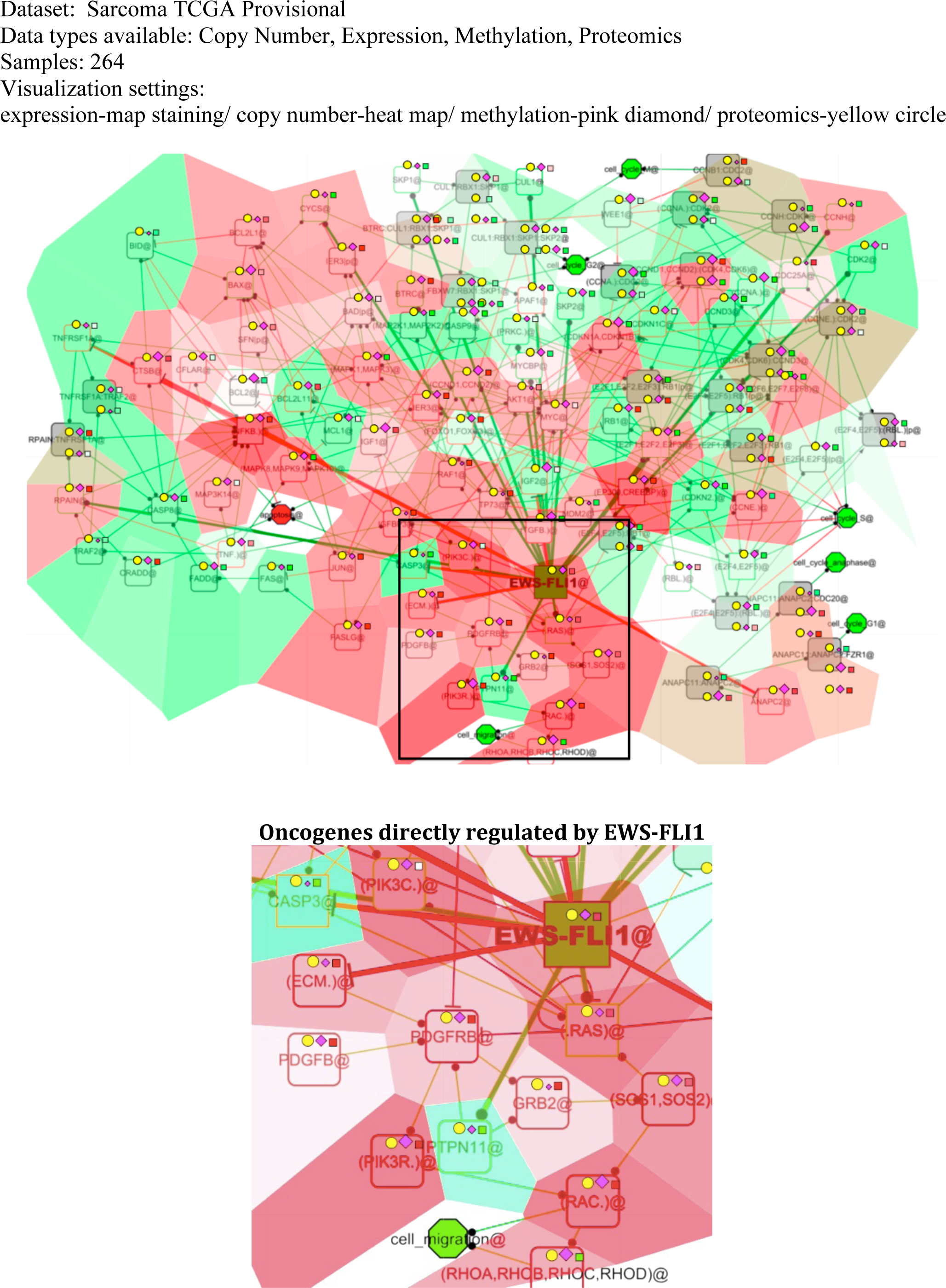
Molecular portrait of Sarcoma visualized on Ewing’s sarcoma map.

**Supplementary figure 3.**
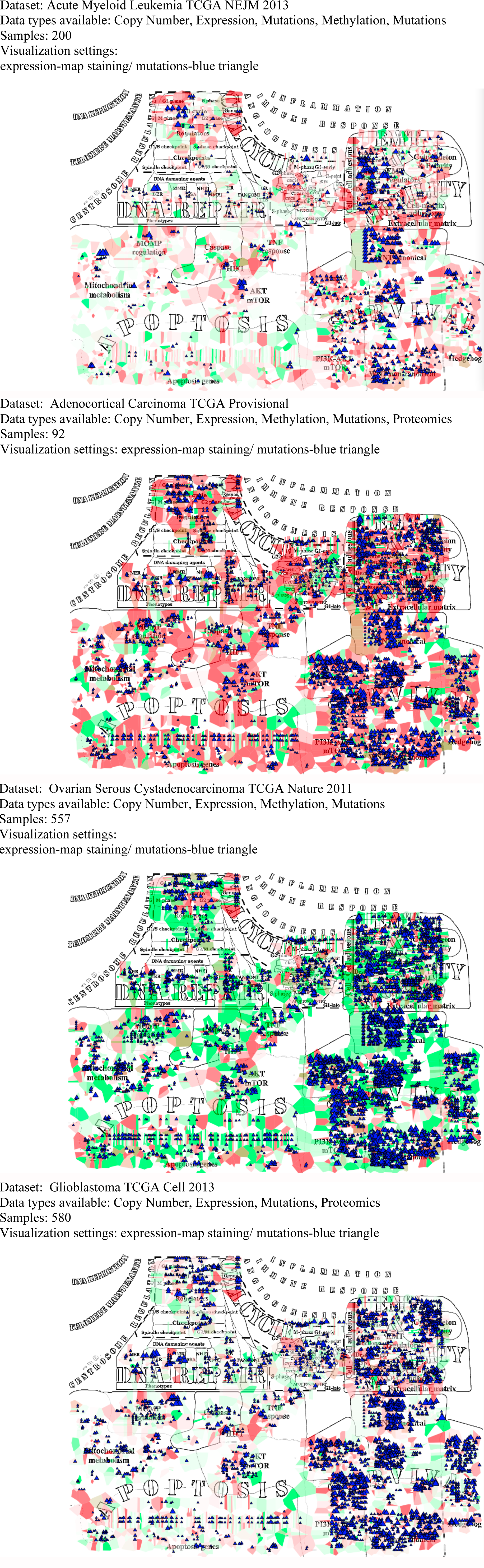
Molecular portrait of several cancer types visualized on ACSN map.

## Reference

1. Yuan,Y., Van Allen,E.M., Omberg,L., Wagle,N., Amin-Mansour,A., Sokolov,A., Byers,L.A., Xu,Y., Hess,K.R., Diao,L., et al. (2014) Assessing the clinical utility of cancer genomic and proteomic data across tumor types. Nat Biotechnol, 32, 644–652.

2. Alyass,A., Turcotte,M. and Meyre,D. (2015) From big data analysis to personalized medicine for all: challenges and opportunities. BMC Med Genomics, 8, 33.

3. Cerami,E., Gao,J., Dogrusoz,U., Gross,B.E., Sumer,S.O., Aksoy,B.A., Jacobsen,A., Byrne,C.J., Heuer,M.L., Larsson,E., et al. (2012) The cBio cancer genomics portal: an open platform for exploring multidimensional cancer genomics data. Cancer Discov, 2, 401–4.

4. Lapatas,V., Stefanidakis,M., Jimenez,R.C., Via,A. and Schneider,M.V. (2015) Data integration in biological research: an overview. J Biol Res (Thessalonikē, Greece), 22, 9.

5. Habermann,B., Villaveces,J. and Koti,P. (2015) Tools for visualization and analysis of molecular networks, pathways, and ‐omics data. Adv Appl Bioinforma Chem, 8, 11.

6. Dorel,M., Barillot,E., Zinovyev,A. and Kuperstein,I. (2015) Network-based approaches for drug response prediction and targeted therapy development in cancer. Biochem Biophys Res Commun, 464, 386–91.

7. Fabregat,A., Sidiropoulos,K., Garapati,P., Gillespie,M., Hausmann,K., Haw,R., Jassal,B., Jupe,S., Korninger,F., McKay,S., et al. (2015) The Reactome pathway Knowledgebase. Nucleic Acids Res, 44, D481–7.

8. Kanehisa,M. (2002) The KEGG database. Novartis Found Symp, 247, 91–101; discussion 101–3, 119–28, 244–52.

9. Paz,A., Brownstein,Z., Ber,Y., Bialik,S., David,E., Sagir,D., Ulitsky,I., Elkon,R., Kimchi,A., Avraham,K.B., et al. (2011) SPIKE: a database of highly curated human signaling pathways. Nucleic Acids Res, 39, D793–9.

10. Cerami,E.G., Gross,B.E., Demir,E., Rodchenkov,I., Babur,O., Anwar,N., Schultz,N., Bader,G.D. and Sander,C. (2011) Pathway Commons, a web resource for biological pathway data. Nucleic Acids Res, 39, D685–90.

11. Kamburov,A., Stelzl,U., Lehrach,H. and Herwig,R. (2013) The ConsensusPathDB interaction database: 2013 update. Nucleic Acids Res, 41, D793–800.

12. Perfetto,L., Briganti,L., Calderone,A., Perpetuini,A.C., Iannuccelli,M., Langone,F., Licata,L., Marinkovic,M., Mattioni,A., Pavlidou,T., et al. (2016) SIGNOR: a database of causal relationships between biological entities. Nucleic Acids Res, 44, D548–54.

13. Kuperstein,I., Bonnet,E., Nguyen,H.-A., Cohen,D., Viara,E., Grieco,L., Fourquet,S., Calzone,L., Russo,C., Kondratova,M., et al. (2015) Atlas of Cancer Signalling Network: a systems biology resource for integrative analysis of cancer data with Google Maps. Oncogenesis, 4, e160.

14. Chowdhury,S. and Sarkar,R.R. (2015) Comparison of human cell signaling pathway databases‐‐ evolution, drawbacks and challenges. Database, 2015, bau126–bau126.

15. Yamada,T., Letunic,I., Okuda,S., Kanehisa,M. and Bork,P. (2011) iPath2.0: interactive pathway explorer. Nucleic Acids Res, 39, W412–5.

16. Caspi,R., Altman,T., Billington,R., Dreher,K., Foerster,H., Fulcher,C.A., Holland,T.A., Keseler,I.M., Kothari,A., Kubo,A., et al. (2014) The MetaCyc database of metabolic pathways and enzymes and the BioCyc collection of Pathway/Genome Databases. Nucleic Acids Res, 42, D459–71.

17. Kono,N., Arakawa,K., Ogawa,R., Kido,N., Oshita,K., Ikegami,K., Tamaki,S. and Tomita,M. (2009) Pathway projector: web-based zoomable pathway browser using KEGG atlas and Google Maps API. PLoS One, 4, e7710.

18. Shannon,P., Markiel,A., Ozier,O., Baliga,N.S., Wang,J.T., Ramage,D., Amin,N., Schwikowski,B. and Ideker,T. (2003) Cytoscape: a software environment for integrated models of biomolecular interaction networks. Genome Res, 13, 2498–504.

19. Dogrusoz,U., Erson,E.Z., Giral,E., Demir,E., Babur,O., Cetintas,A. and Colak,R. (2006) PATIKAweb: a Web interface for analyzing biological pathways through advanced querying and visualization. Bioinformatics, 22, 374–5.

20. Bonnet,E., Viara,E., Kuperstein,I., Calzone,L., Cohen,D.P.A., Barillot,E. and Zinovyev,A. (2015) NaviCell Web Service for network-based data visualization. Nucleic Acids Res, 10.1093/nar/gkv450.

21. Uchiyama,T., Irie,M., Mori,H., Kurokawa,K. and Yamada,T. (2015) FuncTree: Functional Analysis and Visualization for Large-Scale Omics Data. PLoS One, 10, e0126967.

22. Kuperstein,I., Cohen,D.P.A., Pook,S., Viara,E., Calzone,L., Barillot,E. and Zinovyev,A. (2013) NaviCell: a web-based environment for navigation, curation and maintenance of large molecular interaction maps. BMC Syst Biol, 7, 100.

23. Perou,C.M., Sørlie,T., Eisen,M.B., van de Rijn,M., Jeffrey,S.S., Rees,C.A., Pollack,J.R., Ross,D.T., Johnsen,H., Akslen,L.A., et al. (2000) Molecular portraits of human breast tumours. Nature, 406, 747–52.

24. Ciriello,G., Gatza,M.L., Beck,A.H., Wilkerson,M.D., Rhie,S.K., Pastore,A., Zhang,H., McLellan,M., Yau,C., Kandoth,C., et al. (2015) Comprehensive Molecular Portraits of Invasive Lobular Breast Cancer. Cell, 163, 506–519.

25. Hanahan,D. and Weinberg,R.A. (2011) Hallmarks of cancer: the next generation. Cell, 144, 646–74.

26. Kitano,H., Funahashi,A., Matsuoka,Y. and Oda,K. (2005) Using process diagrams for the graphical representation of biological networks. Nat Biotechnol, 23, 961–966.

27. Ogishima,S., Mizuno,S., Kikuchi,M., Miyashita,A., Kuwano,R., Tanaka,H. and Nakaya,J. (2016) AlzPathway, an Updated Map of Curated Signaling Pathways: Towards Deciphering Alzheimer’s Disease Pathogenesis. Methods Mol Biol, 1303, 423–32.

28. Stoll,G., Surdez,D., Tirode,F., Laud,K., Barillot,E., Zinovyev,A. and Delattre,O. (2013) Systems biology of Ewing sarcoma: a network model of EWS-FLI1 effect on proliferation and apoptosis. Nucleic Acids Res, 41, 8853–71.

29. Bonnet,E., Calzone,L., Rovera,D., Stoll,G., Barillot,E. and Zinovyev,A. (2013) BiNoM 2.0, a Cytoscape plugin for accessing and analyzing pathways using standard systems biology formats. BMC Syst Biol, 7, 18.

30. Uchiyama,T., Irie,M., Mori,H., Kurokawa,K. and Yamada,T. (2015) FuncTree: Functional Analysis and Visualization for Large-Scale Omics Data. PLoS One, 10, e0126967.

31. Yamada,T., Letunic,I., Okuda,S., Kanehisa,M. and Bork,P. (2011) iPath2.0: interactive pathway explorer. Nucleic Acids Res, 39, W412–5.

32. Kono,N., Arakawa,K., Ogawa,R., Kido,N., Oshita,K., Ikegami,K., Tamaki,S. and Tomita,M. (2009) Pathway projector: web-based zoomable pathway browser using KEGG atlas and Google Maps API. PLoS One, 4, e7710.

33. Croft,D., Mundo,A.F., Haw,R., Milacic,M., Weiser,J., Wu,G., Caudy,M., Garapati,P., Gillespie,M., Kamdar,M.R., et al. (2014) The Reactome pathway knowledgebase. Nucleic Acids Res, 42, D472–7.

34. Kanehisa,M., Goto,S., Sato,Y., Furumichi,M. and Tanabe,M. (2012) KEGG for integration and interpretation of large-scale molecular data sets. Nucleic Acids Res, 40, D109–14.

35. Croft,D., O’Kelly,G., Wu,G., Haw,R., Gillespie,M., Matthews,L., Caudy,M., Garapati,P., Gopinath,G., Jassal,B., et al. (2010) Reactome: a database of reactions, pathways and biological processes. Nucleic Acids Res, 39, D691–7.

